# Failure of digit tip regeneration in the absence of *Lmx1b* suggests Lmx1b functions disparate from dorsoventral polarity

**DOI:** 10.1101/2022.05.05.490742

**Authors:** Alejandro Castilla-Ibeas, Sofía Zdral, Laura Galán, Endika Haro, Lila Allou, Víctor M. Campa, Jose M. Icardo, Stefan Mundlos, Kerby C. Oberg, Marian A. Ros

## Abstract

Mammalian digit tip regeneration is linked to the presence of nail tissue, but a nail-explicit model is missing. Here, we report that nail-less double-ventral digits of Δ*LARM1/2* mutants that lack limb-specific *Lmx1b* enhancers fail to regenerate. To separate the nail’s effect from the lack of DV polarity, we also interrogate double-dorsal double-nail digits and show that they regenerate. Thus, DV polarity is not a prerequisite for regeneration and the nail requirement is supported. Transcriptomic comparison between wild-type and non-regenerative *ΔLARM1/2* mutant blastemas reveals differential up-regulation of vascularization and connective tissue functional signatures in wild-type versus upregulation of inflammation in the mutant. These results, together with the finding of uniform *Lmx1b* expression in the wild-type blastema and in the dorsal dermis underneath the nail, indicate that, in addition of the nail’s effect, a direct role for Lmx1b in driving the progression of digit tip regeneration is likely.

## INTRODUCTION

The ability to regenerate damaged or lost tissues varies greatly across the animal kingdom. One type of regenerative response, known as epimorphic regeneration, restores the original anatomy and function of amputated structures. The process depends on the formation of a blastema, a transient structure composed of morphologically similar undifferentiated cells that form at the end of the amputated stump under the wound epithelium. The blastema cells proliferate and eventually differentiate to form the regenerate that faithfully recapitulates the form and function of the lost structure. A dramatic example of epimorphic regeneration is exhibited by salamanders with the capacity to regrow a complete limb if amputated (Bassat and Tanaka 2021). In contrast, mammals have a very limited capacity for epimorphic regeneration as wound healing normally resolves with the formation of a fibrotic non-functional scar at the wound site (Simkin et al. 2013). However, one notable exception is the ability of rodents and primates (including humans) to regenerate digit tips (Borgens 1982; Illingworth 1974). The regeneration of a digit tip replicates many aspects of the epimorphic regeneration seen in other organisms including the formation of a blastema, scarless wound healing, and regrowth of the lost part with essentially the same form and function as what was amputated (Suzuki et al. 2006; Johnson and Lehoczky 2022; Stocum 2017).

The regeneration of the digit tip has been functionally associated with the presence of the nail/claw because only amputations distal to (or that preserve) the nail matrix regenerate (Zhao and Neufeld 1995; Han et al. 2008; Takeo et al. 2013). The nail is a dorsal keratinized ectodermal derivative whose continuous growth depends on a population of nail stem cells located in the nail matrix and characterized by the expression of *Lgr6* (Lehoczky and Tabin 2015). The influence of the nail in digit tip regeneration has been attributed to Wnt signaling emanating from the distal nail matrix that promotes nail differentiation. After amputation, Wnt signaling from the distal nail matrix induces nerve-dependent Fgf2 secretion from nail epithelium, which in turn activates the underlying blastema (Takeo et al. 2013 and reviewed in Lehoczky 2017). Accordingly, removal of β–catenin in nail epithelium prevents blastemal growth and regeneration in distal amputations, while stabilization of β–catenin extends the regenerative capacity to a more proximal level (Takeo et al. 2013). Nevertheless, a more explicit test of the nail’s role in digit tip regeneration would be from a nail-less digit model.

The nail is a distinctive dorsal feature whose development is linked to limb dorsoventral (DV) patterning. During early limb development, the ectoderm covering the emerging bud is divided into dorsal ectoderm expressing *Wnt7a* and ventral ectoderm expressing *En1* (Tickle 2015). Wnt7a from the dorsal ectoderm induces the expression of the LIM-homeodomain transcription factor Lmx1b in the subjacent mesoderm. *Lmx1b* is transcribed precisely in the limb progenitors located in the dorsal half of the bud where it is necessary and sufficient to specify dorsal fates (Vogel et al. 1995; Chen et al. 1998; Riddle et al. 1995). *Lmx1b* controls dorsal patterning of the skin (integument), of the internal organs (musculoskeletal components) and it also provides specific guidance cues for spinal motor axons (Krawchuk and Kania 2008).

Targeted disruption of *Lmx1b* results in limbs that are double-ventral at zeugopod and autopod level (Chen et al. 1998). Double-ventral (or bi-ventral) limbs refer to limbs whose dorsal half is a mirror image of the ventral side. Thus, the dorsal surface of *Lmx1b-null* limbs adopts the typical ventral appearance with sparse hair and is most striking in hands and feet because of the absence of nails and the presence of ectopic dorsal pads (Chen et al. 1998) *Lmx1b-null* mice would be an excellent model to test the requirement of the nail for digit tip regeneration, but unfortunately, they die perinatally due to multiple systemic defects associated with *Lmx1b* pleiotropy. *Lmx1b* plays essential roles in a wide range of developmental processes including the development of the kidney, the anterior segment of the eye, and central nervous system (specifically dopaminergic and serotonergic neurons) (Sweeney et al. 2003). Recently, we characterized two limb-specific *Lmx1b* enhancers, termed *LARM1* and *LARM2* (*<u>L</u>mx1b* <u>a</u>ssociated <u>r</u>egulatory <u>m</u>odule 1 and 2) that are key regulators of *Lmx1b* expression in the limb (Haro et al. 2017; 2021). Accordingly, deletion of both *LARM* enhancers results in mice (*ΔLARM1/2*) that replicate the *Lmx1b-null* phenotype exclusively in the limb and are, therefore, viable. The double-ventral limbs with non-claw bearing digits of *ΔLARM1/2* mice make them an attractive model to directly interrogate the role of the nail in digit tip regeneration.

Here, we report that *ΔLARM1/2* homozygous mutants fail to regenerate amputated digit tips supporting the nail requirement. However, we also tested for two caveats that could challenge this conclusion. One was the possibility that the absence of polarity in the DV axis of *ΔLARM1/2* mutants could impair regeneration. This notion comes from amphibian studies showing a correlation between positional information disparity and the ability to regenerate (Bryant and Gardiner 1992). To disentangle the absence of polarity from the lack of a nail, we took advantage of the *Del(27)* mutant that also lacks polarity due to the limb-restricted loss of *En1* (Allou et al. 2021) and displays ectopic ventral *Wnt7a/Lmx1b* expression with circumferential nails. We found that *Del(27)* digits do regenerate, indicating that asymmetrical DV positional information is not required for regeneration. The second caveat comes from the finding that, in the wild-type, *Lmx1b* expression persisted into adulthood, detected in the connective tissue underneath the nail, the onychodermis. Interestingly, *Lmx1b* is also expressed at low level and without DV bias in the whole WT blastema raising the possibility that *Lmx1b* expression directly contributes to regenerative capacity. In the absence of *Lmx1b*, the regeneration process initiates normally, but stalls after the formation of a blastema. The comparison of the transcriptomic signature of the regenerative control vs the non-regenerative mutant blastema points to impaired vasculogenesis, extracellular matrix (ECM) remodeling and exacerbated inflammation as possibly causal factors. Conjointly, our results suggest a direct influence of Lmx1b to promote regeneration, an view that will require further investigation.

## RESULTS

### *ΔLARM1/2* mutants display digit tips with double-ventral morphology

We have recently characterized two limb-specific *Lmx1b* enhancers, termed *LARM1* and *LARM2*, that mediate Lmx1b limb dorsalization (Haro et al. 2021). Mice homozygous for the removal of these two enhancers (*ΔLARM1/2*) replicate the *Lmx1b-null* limb phenotype consisting of double-ventral distal limbs with sole pads in both surfaces and no nails (Haro et al. 2021; Fig 1A-A’). In contrast to *Lmx1b-null* mutants that die perinatally, *ΔLARM1/2* homozygous mutants survive, as they show no other Lmx1b-related systemic defects, allowing the analysis of fully differentiated double-ventral limbs. Visual inspection of the dorsal surface of the 3-week-old wild type (WT) and mutant paws showed almost complete absence of hair and the presence of typical ventral features such as volar pads and nail agenesis in *ΔLARM1/2* homozygous mutants (Fig. 1A-A’). Lateral examinations of the digit tips, both grossly (Fig. 1B-B’) or under scanning electron microscopy (Fig. 1C-C’) confirmed the absence of the nail organ that was replaced by an ectopic dorsal pad in the mutant. The lateral views also exposed the DV symmetry of the mutant digit tip while the histological analysis of trichrome stained longitudinal sections revealed the DV symmetry of the skeletal elements (Fig. 1D-D’). The morphology of the distal phalanx was consistent with the apposition of two ventral halves, the dorsal extensor tendon adopted the mirror image shape of the ventral flexor tendon and the structure of the ectopic dorsal digital pad (DDP) mirrored that of the volar digital pad with the inclusion of eccrine glands (Fig. 1D-D’). Corresponding schemes are shown in Fig. 1E-E’.

**Figure 1.**
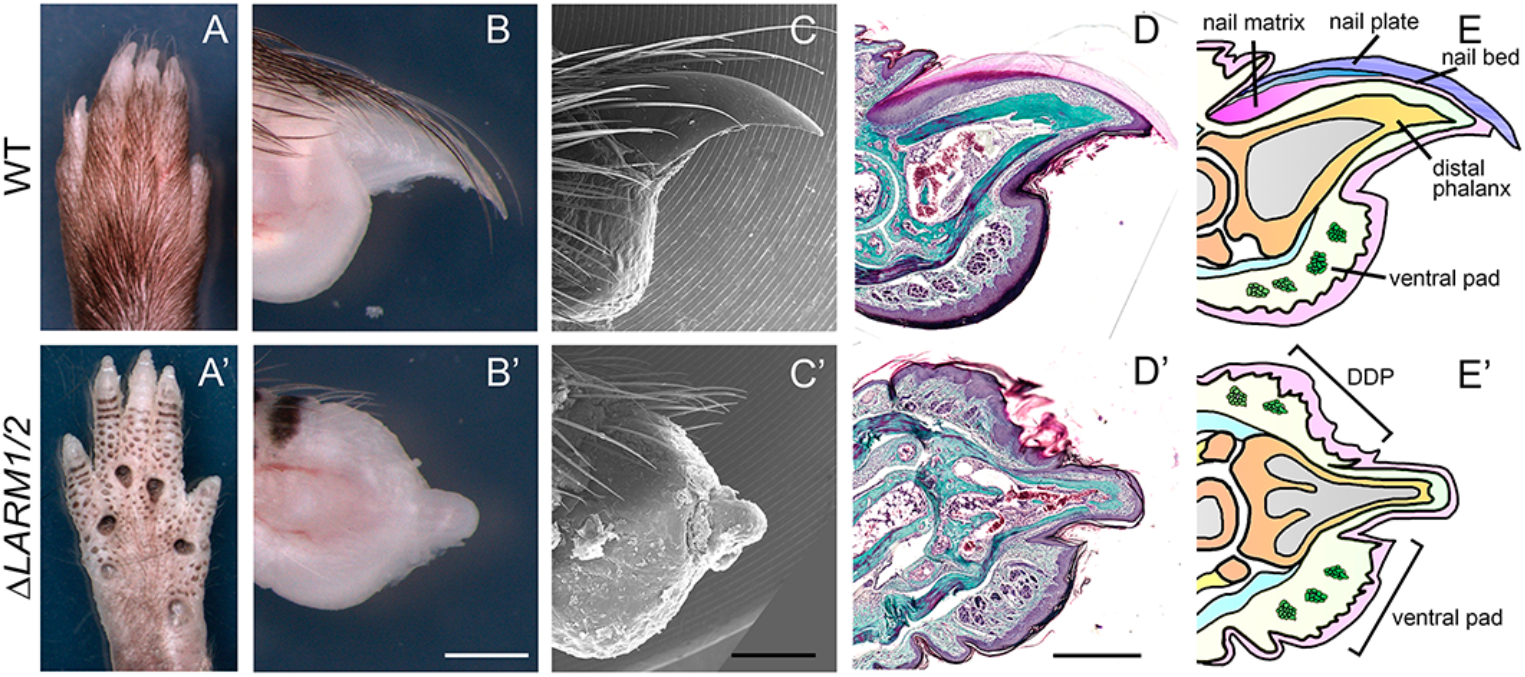
*ΔLARM1/2* mutants display digits with double-ventral morphology. (A-A’) Dorsal gross morphology of the hindlimb autopod of WT (A) and *ΔLARM1/2* (A’) mice. (B-B’) Lateral gross morphology of a hindlimb digit tip of WT (B) and *ΔLARM1/2* (B’) mice. (C-C’) Scanning electron microscopy microphotographs of P21 digit tips from WT (C) and *ΔLARM1,2* (C’) mice. (D-D’) Masson’s trichrome staining of P21 longitudinal sections from WT (D) and *ΔLARM1/2* (D’) digit tips. (E-E’) Schematics of the digit tip of WT (E) and *ΔLARM1/2* (E’) mice. DDP: dorsal digital pad Scale bars: 500 μm

Over time the DDP developed several epidermal invaginations that were initially unnoticeable from the surface (Fig. 1B’-C’), but eventually produced a highly keratinized structure. Because these structures formed in place of the nail, and our goal was to understand the role of the nail in digit tip regeneration, we decided to perform an in-depth study of the DDP at both morphological and molecular levels.

### The dorsal tips of *ΔLARM1/2* homozygous digits form a hybrid nail-pad structure

To evaluate the nature of the DDP formed in the mutant nail region, we examined its epithelial differentiation using a range of molecular markers either by immunohistochemistry (IHC) or by in situ hybridization (ISH). We checked the expression of Keratin (Krt) 5 and Krt10 markers of the basal and supra-basal skin layers, respectively. The normal differentiation of the nail matrix during the perinatal period is characterized by expression of Krt5 in the basal layer of the epidermis, specific downregulation of Krt10 in the suprabasal layers, and the activation of hard keratins specific for hairs and nails, as detected with the AE13 antibody, at postnatal day (P) 21 (Fig. 2A-C; Fernandez-Guerrero et al. 2020). In *ΔLARM1/2* homozygous mutants, the epidermal differentiation of the DDP followed the nail program, although attenuated, as Krt10 was downregulated concomitantly with the expression of low levels of hard keratins specifically in the epithelial invaginations of the DDP (Fig. 2A’-C’).

**Figure 2.**
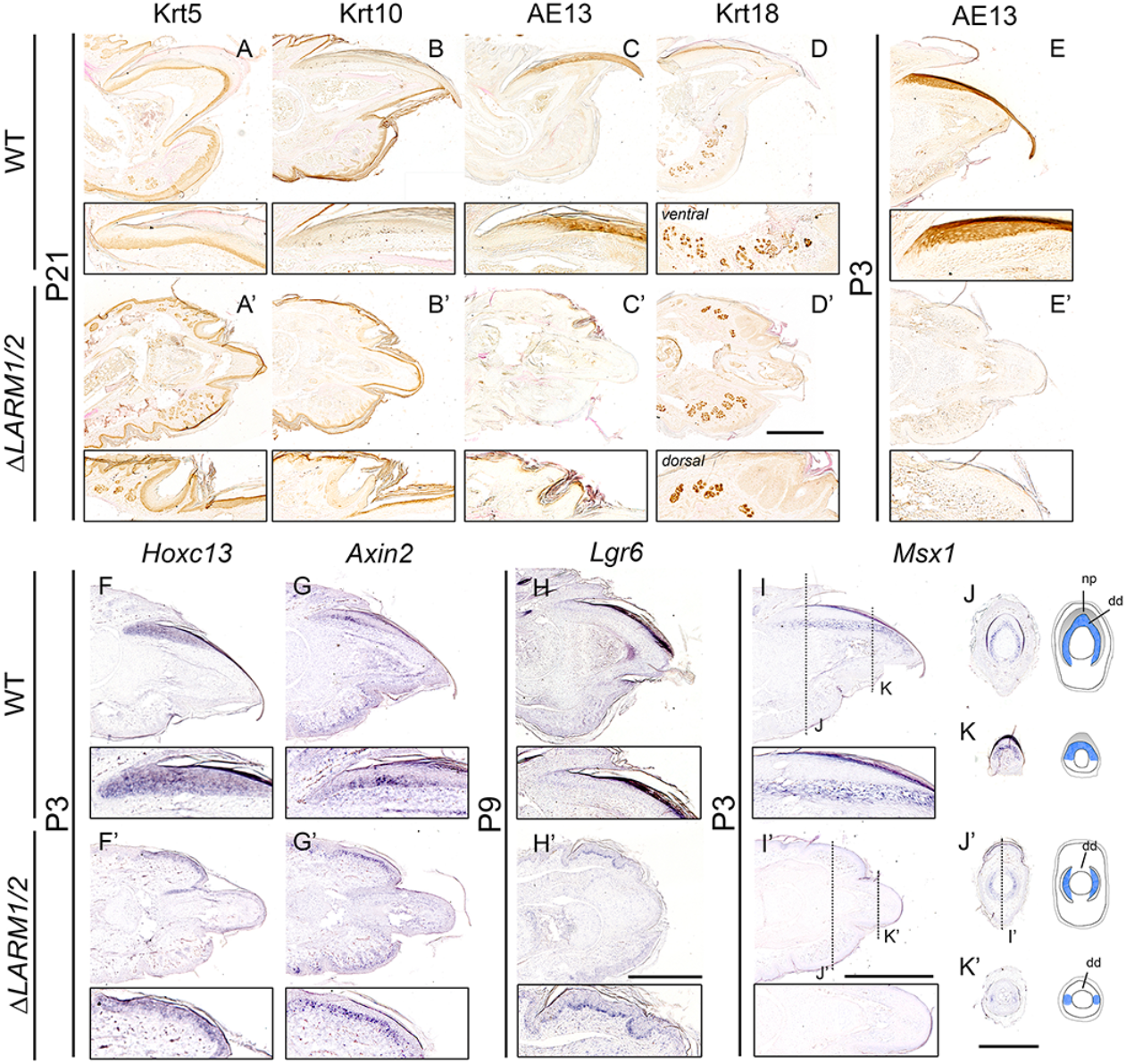
The dorsal tip of *ΔLARM1/2* homozygous digits form a hybrid nail-pad structure. (A-E’) Immunohistochemistry for the detection of Krt5 (n=5), Krt10 (n=4), Krt18 (n=3) and hard keratins (AE13 antibody, n=3 for P21) in equivalent longitudinal sections of P21 and P3 (n=2) digit tips of WT and *ΔLARM1/2* mutants. (F-I’) ISH for *Hoxc13* (n=2), *Axin2* (n=2), *Lgr6* (n=2) and *Msx1* (n=4) in longitudinal sections of P3 or P9 digit tips of WT and *ΔLARM1/2* samples as indicated. (J-K’) Left: transversal sections stained by ISH for *Msx1*, the planes are indicated in H and H’. Right: corresponding schemes depicting the expression in blue. np: nail plate, dd: dorsal dermis The inset below each image in (A-I’) shows a 1.5× magnification of the dorsal area except from D and H’, where the ventral and central portion of the digit is magnified, respectively. Scale bars: 500 μm

We also checked for the presence of sweat glands, a ventral characteristic, in the DDP of *ΔLARM1/2* mutants using the Krt18 marker (Lu et al. 2016). At P21, eccrine sweat glands were prominent in the dorsal pad, although always in lower number than in the ventral pad (arrow Fig. 2D-D’). It should be mentioned that at P3, Krt18 positive eccrine glands were not detected in the mutant DDP, consistent with the reported absence in the DDP of *Lmx1b*-null mice (Chen et al. 1998). Thus, the dorsal side of the *ΔLARM1/2* mutant digit tip displays a structure similar to a ventral pad containing eccrine glands, but with nail-specific epidermal features such as the expression of hard keratins.

To further explore the state of the nail differentiation program in *ΔLARM1/2* mutants, we used ISH to examine the expression of *Hoxc13*, a transcriptional regulator of hair and nail specific hard keratins and a marker of the nail matrix (Fernandez-Guerrero et al. 2020). Surprisingly, low, but clear, *Hoxc13* expression demarcated the epidermis of the DDP already at P3 (Fig. 2F-F’), although not accompanied by the expression of hard keratins at this stage (Fig. 2E-E’). The expression of *Axin2*, a negative regulator and target of the Wnt signaling pathway, was also analyzed as a marker of epidermal stem cells (Lim et al. 2013). In the WT skin, expression of *Axin2* was observed in scattered cells of the basal layer of the interfollicular epidermis and more intensely in the nail matrix coincident with the location of the nail stem cells. In *ΔLARM1/2* mutants, *Axin2* expressing cells were similarly observed in the interfollicular epidermis and more densely in the epidermis of the DDP suggesting the presence of nail stem cells (Fig. 2G-G’). Indeed, those cells also expressed *Lgr6* (Fig. 2H-H’), considered the best marker of nail stem cells (Lehoczky and Tabin 2015). Finally, we also examined the expression of *Msx1* (Fig. 2I-K’), a marker of the dorsal dermis underneath the nail (Reginelli et al. 1995; Han et al. 2008), the equivalent to the region called onychodermis in humans (D. Y. Lee et al. 2012; Sellheyer and Nelson 2013), a specialized connective tissue region implicated in digit tip regeneration (Lehoczky, Robert, and Tabin 2011; Storer et al. 2020). In *ΔLARM1/2* mutants, the dorsal dermis was significantly reduced in size, equivalent to the ventral dermis in controls, and *Msx1* expression was nearly absent in this region, mimicking the very low level in the ventral dermis (Fig. 2I-I’). Interestingly, *Msx1* expression remained in the lateral dermis best seen in transverse sections (J-K’ and accompanied scheme). This indicates that the dorsal expression of *Msx1* demarcates a unique dorsal domain likely dependent on *Lmx1b*.

Thus, we conclude that the DDP that forms at the tip of *ΔLARM1/2* digits display nail-specific characteristics such as the enrichment of stem cells marked by the expression of *Hoxc13, Axin2* and *Lgr6*. This is consistent with an attenuated nail-like type of epidermal differentiation initially spanning the whole DDP and later concentrating in the invaginations of the pad. The enrichment of stem cells in these invaginations correlates with the high level of keratinization in this structure. However, the dorsal dermis is markedly reduced in size and with no evidence of its defining gene expression (*Msx1*).

### No detectable *Lmx1b* expression in *ΔLARM1/2* double-ventral digits

Given the presence of some, previously unnoticed, nail features in the DDP of *ΔLARM1/2* digits and since the nail is a dorsal structure considered to be dependent on Lmx1b for differentiation, we decided to carefully examine the expression of *Lmx1b* in the *ΔLARM1/2* mutant digits that do not carry a constitutive deletion of *Lmx1b* but of its limb-specific enhancers. To this end, we performed a comprehensive analysis of *Lmx1b* expression in *ΔLARM1/2* mutants using RNA-FISH with amplification by hybridization chain reaction (HCR) that increases sensitivity but also allows quantification and pattern examination (Choi et al. 2018) (Fig. 3). Because *Lmx1b* expression persists into the perinatal period (Schweizer, Johnson, and Brand-Saberi 2004), our analysis spanned embryonic and postnatal stages to capture the dynamics of *Lmx1b* expression in mouse digits. During normal limb development, robust *Lmx1b* expression was found restricted to the dorsal mesoderm (as shown for E11.5 in Fig. 3A). Shortly after birth (P3), *Lmx1b* expression persisted, albeit reduced, in the dorsal dermis of control digits, immediately below the nail organ in an area coincident with the onychodermis (Fig. 3B). At 3 weeks of age (P21), *Lmx1b* expression was further reduced, but still detectable in the dorsal dermis (Fig. 3C-E). In contrast, no *Lmx1b* expression was detected in the digits of *ΔLARM1/2* mutants during development (E11.5, Fig. 3A’) nor at any later stage analyzed (Fig. 3B’ and Fig. 3C, D, F). Quantification of the HCR-FISH signals at P21 performed by counting the number of fluorescent puncta in identical-sized square regions confirmed no significant differences between expression in the mutant dorsal dermis and background levels (absence of initiator probe) while significant differences were found with *Lmx1b* expression in the WT dorsal dermis (Fig. 3C-D).

**Figure 3.**
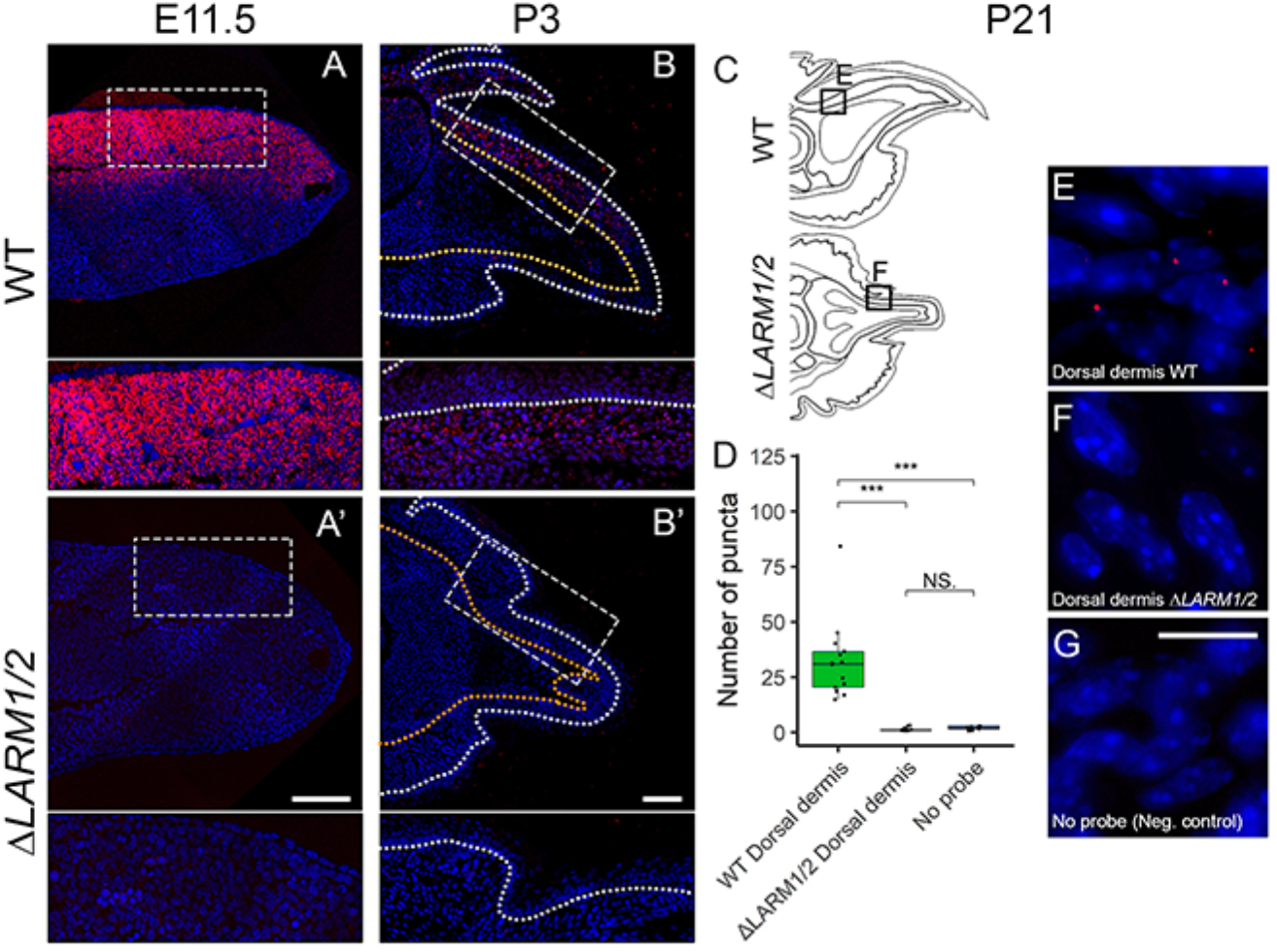
*Lmx1b* expression is not detectable in *ΔLARM1/2* mutants. (A-A’) HCR RNA-FISH for *Lmx1b* in longitudinal sections of E11.5 control and *ΔLARM1/2* mutants hindlimb buds (n=3) (B-B’) HCR RNA-FISH for *Lmx1b* in longitudinal sections of P3. White and orange dotted lines mark the epidermal-dermal boundary and distal phalanx bone, respectively (n=3). Scale bars represent 50 μm. Blue: DAPI Insets below A-B’ panels show a 2x magnification of the framed region. (C) Schematics of WT and *ΔLARM1/2* P21 digit tips marking the studied areas. (D) Boxplot of the number of *Lmx1b* puncta detected by HCR-FISH at P21 in the WT dorsal dermis, *ΔLARM1/2* dorsal dermis, and in no-probe negative control sections (n=3). Each dot is a measurement in different images from the same regions. (E-F) Representative images of the regions quantified in D as indicated in the schematics in C. (G) Representative image of a section incubated without probes as a negative control. Scale bar: 10 μm Blue: DAPI

Therefore, in view of our analysis, we concluded that part of the differentiation program of the nail organ occurs in *ΔLARM1/2* mutants in the complete absence of detectable *Lmx1b* expression. It should be noted that the limited residual dorsal features of *ΔLARM1/2* homozygous were exclusively detected in the ectoderm, while the morphology of the internal elements (bone and tendons) was totally double-ventral (Fig. 1 and 2). Moreover, characteristic dorsal gene expression in the nail dermis (i.e., *Msx1*) was absent.

### The bi-ventral digits of *ΔLARM1/2* mutants fail to regenerate

The presence of the nail/claw is considered a requirement for digit tip regeneration (Zhao and Neufeld 1995; Takeo et al. 2013; Lehoczky and Tabin 2015). Although we show that some aspects of the nail differentiation program are present in *ΔLARM1/2* mutant ectoderm, a true nail organ is never established, neither is a nail plate. Therefore, we asked whether *ΔLARM1/2* mutants could regenerate after permissive amputations (those that eliminate the distal third of the distal phalanx) in three-week-old (P21) mice and monitored the progressive regeneration process. In high contrast to WT mice, *ΔLARM1/2* mutants did not regenerate their digit tips as assessed by micro-computed tomography (μCT) 28 days post amputation (dpa) (n=5; Fig. 4A-C’ and G). As previously described, at this time point regeneration was considered completed except for the final bone remodeling (Fernando et al. 2011). Because the regenerative ability decreases over time (Brunauer et al. 2021; Yun 2015), we also investigated whether younger *ΔLARM1/2* mutants were capable of regeneration by testing the amputation response in three-day-old (P3) animals (n=4; Supplemental Fig. 1). Our results showed that permissive amputations in mutant digits at P3 also failed to regenerate. Therefore, the DDP that forms in *ΔLARM1/2* mutants, despite its nail-like features, is not sufficient for regeneration. This finding supports either the notion that a completely well-formed nail organ is required for regeneration or that other mechanisms are impeding regeneration in the mutant (see below).

**Figure 4.**
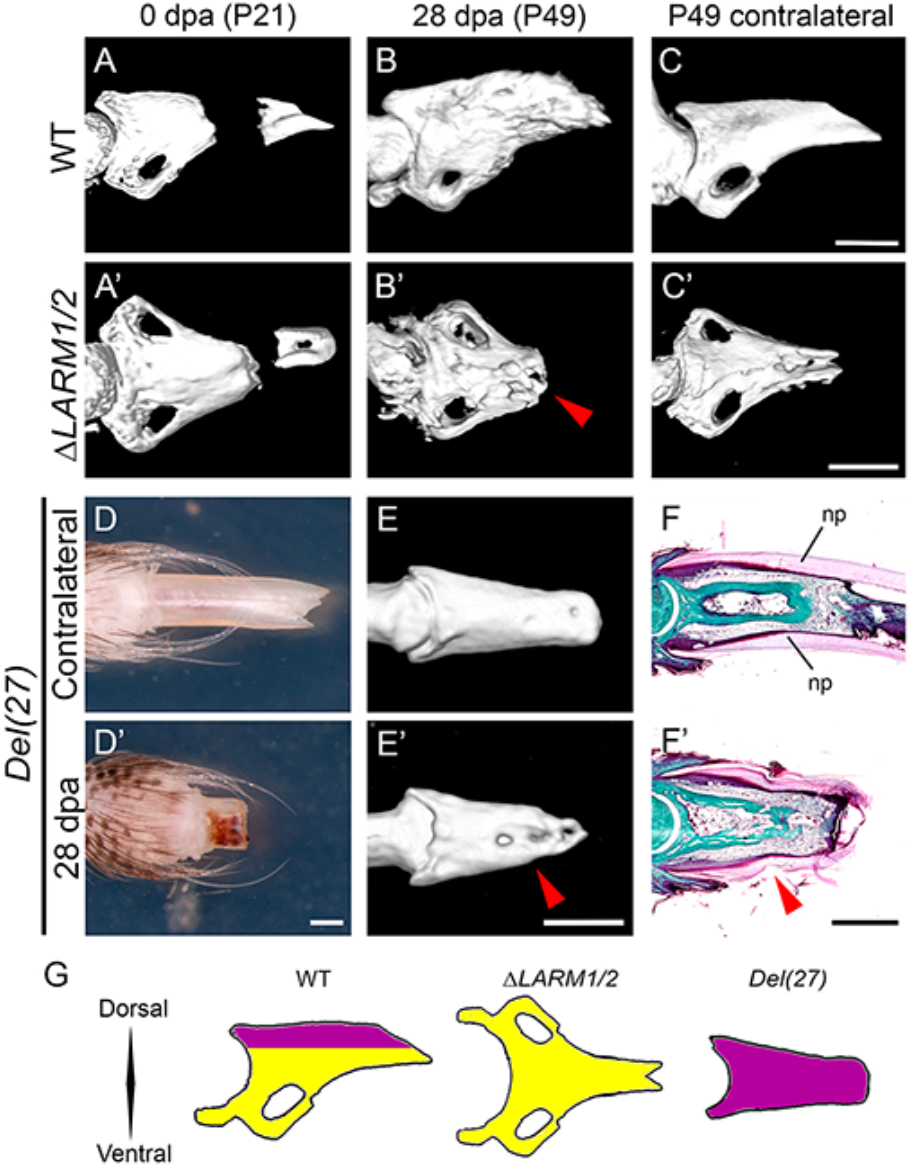
*ΔLARM1/2* mutants do not regenerate amputated digit tips but digit tip regeneration occurs in the absence of DV polarity. (A-C’) μCT scans of P21 WT and *ΔLARM1/2* digit tips immediately after amputation showing the amputated tip (A-A’) and 28 dpa (B-B’) with the unamputated contralateral digit tips (P49) as controls (C-C’) (n=5). (D-D’) Lateral gross morphology of a *Del(27)* mutant digit tip 28 dpa (D’) and the unamputated contralateral digit (P49) as control (D). (E-E’) μCT scans of a *Del(27)* mutant digit tip 28 dpa (E’) and unamputated contralateral digit (P49) as control (E). (F-F’) Masson’s trichrome staining of a longitudinal section of a *Del(27)* mutant digit tip at P49, unamputated (F) and 28 dpa (F’). (G) Schematics depicting the morphology of the distal phalanx of WT, double ventral and double dorsal digits. The dorsal part is colored in purple and the ventral part in yellow. Scale bars: 500 μm

### The bi-dorsal digits of *Del(27)* mutants do regenerate despite lack of DV polarity

We first considered the possibility that the lack of DV polarity in *ΔLARM1/2* mutants, rather than the lack of a complete nail organ, impairs the ability to regenerate the digit tip. This concept is supported by classical experiments in amphibians showing that double-ventral or double-dorsal limbs failed to elicit normal distal regeneration (Bryant and Baca 1978; Satoh and Makanae 2014), although conflicting results have also been reported (Burton, Holder, and Jesani 1986). We reasoned that if DV polarity is a prerequisite for regeneration, a double-dorsal limb, with digits bearing conical or circumferential nails, would not regenerate, despite an ample nail organ. To test this hypothesis, we used the recently generated *Del(27)* mutant that because of the loss of the *Maenli* lncRNA, a master activator of *En1* in the limb, displays a double-dorsal limb phenotype with digits carrying both dorsal and ventral nails (Allou et al. 2021; Fig. 4D-G). We found that *Del(27)* mutants did regenerate their digit tips after amputation at P21 (Fig. 4D’-F’), demonstrating that digits without DV polarity are capable of regenerating. Moreover, this finding further supports the role of a nail organ as a necessary component of digit tip regeneration.

### Despite regeneration failure, a blastema forms in the amputated stump of *ΔLARM1/2* mutant digits

Searching for the reason for the lack of regeneration in the *ΔLARM1/2* mutant, we monitored the regeneration process in histologic sections after amputations at P21 (Supplemental Fig. 2A-F’). Histolysis is an important step in the initiation of digit tip regeneration characterized by the enzymatic degradation of damaged tissue and correlates with the size of the blastema (Simkin et al. 2015). Osteoclast-mediated histolysis of the distal bone after an amputation is thought to facilitate regeneration by releasing extracellular matrix-linked mitogens/chemokines and exposing bone marrow precursors and mitogens to the amputation site (Simkin et al. 2015). The histolysis response is clearly present in *ΔLARM1/2* digits, demonstrated in trichrome stains by the ejection of distal bone fragments external to the wound epithelium, still embedded in residual clot, and by the presence of osteoclasts (Supplementary Fig. 2B’). Additionally, in situ hybridization for *Ctsk*, an osteoclast marker, revealed comparable populations of osteoclasts in WT and mutant digit tips at 7dpa (Fig 5A-A’).

**Figure 5.**
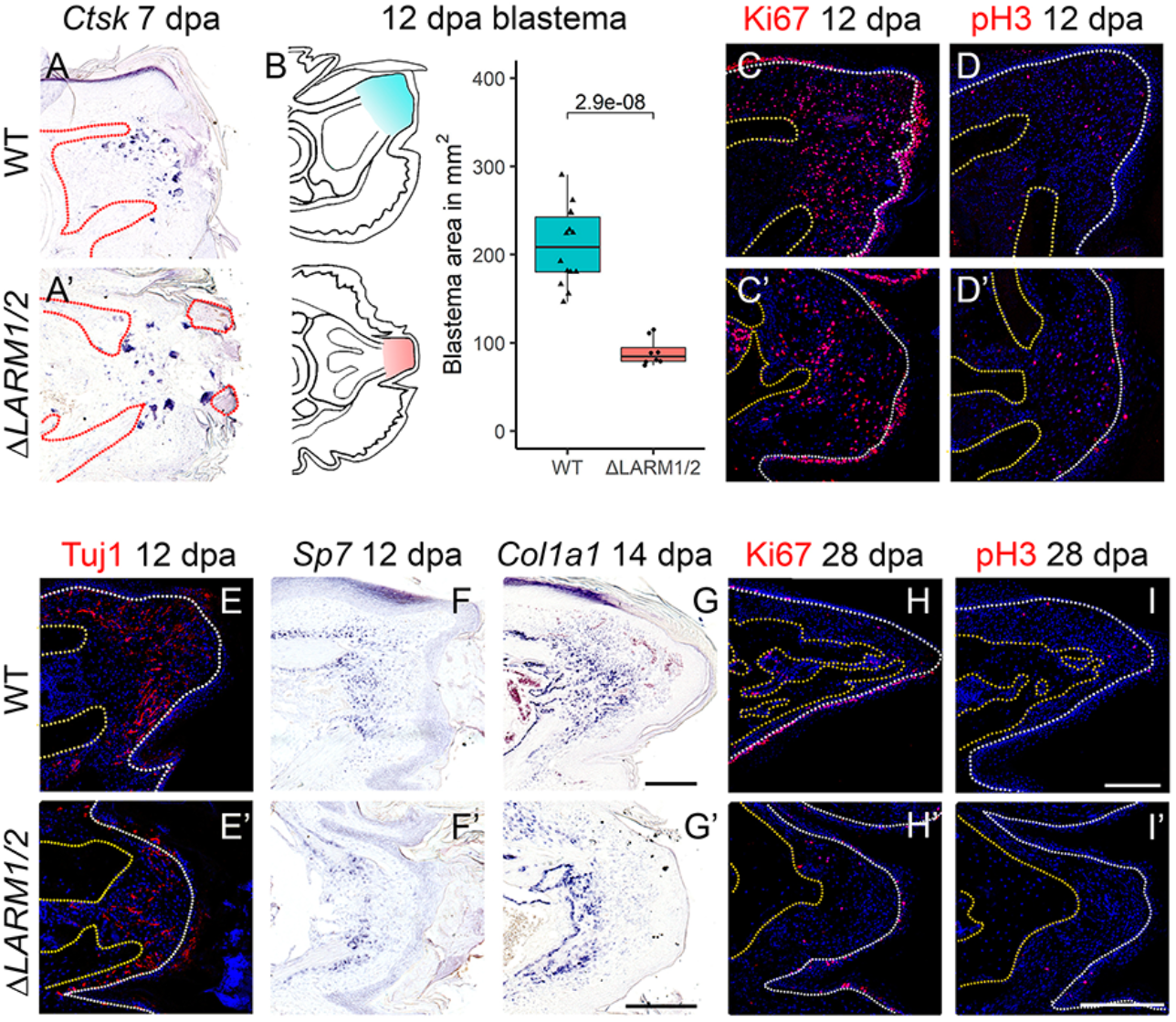
Molecular features of the Δ*LARM1/2* non-regenerative blastema. (A-A’) ISH in longitudinal sections with probes for *Ctsk* of WT (A) and *ΔLARM1/2* (A’) digit tips amputated at P21 (n=2). (B) Left, schematics indicating the region where both blastemas are located, right, quantification of the area in mm^2^ in central sections of 12 dpa blastemas, significance assessed by Student’s t-test (n=4). (C-E’) Immunofluorescence for the detection of Ki67 (n=3), pH3 (n=2) and Tuj1 (n=3) (red) in equivalent longitudinal sections of WT (C,D,E) and *ΔLARM1/2* (C’,D’,E’) digit tips amputated at P21. (F-G’) ISH in longitudinal sections with probes for *Col1a1* (n=3) and *Sp7* (n=2) as indicated on top of WT (F,G) and *ΔLARM1/2* (F’,G’) digit tips amputated at P21. (H-I’) IF for Ki67 (n=2) and pH3 (n=2) at 28 dpa of WT (H,I) and *ΔLARM1/2* (H’,I’) digit tips amputated at P21. Days post-amputation is indicated on top. Scale bars: 500 μm for ISH, 200 μm for IF. White and red/orange dotted lines outline the epidermal and bone boundaries, respectively. Blue in IF: DAPI counterstain.

Most interestingly, despite the absence of regeneration, a blastema forms between the proximal bone stump and the distal wound epidermis (WE) in mutant digits (Supplemental Fig. 2). At a histologic level, the mutant blastemas had the same appearance as the control blastemas; however, they were significantly smaller (Fig. 5B; 12 dpa; p=2.9e-08), and eventually failed to differentiate to form the missing distal end of the phalanx.

At present, there is not a specific marker for blastemal cells, but several features have been described that can help in their identification (Seifert and Muneoka 2018). First, we monitored the proliferative state of blastema cells using immunofluorescence (IF) for Ki67, a marker for cell proliferation status, and pH3, a marker for the late G2 and mitosis stages of the cell cycle. At 12 dpa, Ki67 positive and pH3 positive cells were conspicuous along the time course of blastema formation and development both in control and mutant, but more irregularly distributed in the mutant (Fig. 5C-D’). At this time point, we also checked for nerve colonization, a recognized characteristic of regenerative blastemas (Takeo et al. 2013; Rinkevich et al. 2014; Johnston et al. 2016). IF with the neuron-specific anti-beta-III tubulin antibody (TuJ-1) detected the presence of axon cells in the mutant blastemas, although less abundant than in control blastemas (Fig. 5 E-E’).

Eventually, digit tip regeneration relies on the differentiation of blastema cells to reform the original cell types. Restoration of the amputated distal phalanx is mediated by direct ossification from blastema cells that express osteoblast markers such as *Sp7/Osx* (Dawson et al. 2018). Expression of *Sp7* at 12 dpa marked osteogenic precursors in WT extending from the periosteal and endosteal regions of the bone stump into the blastema while in the mutant, *Sp7* positive cells were found restricted to the bone stump but not in the blastema (Fig. 5 F, F’). The analysis of *Col1a1*, another osteoblast marker, at later blastema stages (14 dpa) confirmed that osteoblasts were largely restricted to the distal end of the phalanx near the site of amputation in mutants while conspicuous in the WT blastema (Fig. 5G-G’). Thus, although some minor bone formation may occur at the tip of the amputation stump in mutants, no osteogenic differentiation is detected in the non-regenerative blastema, in accordance with the μCT results (compare Fig 4B’ with 4C’). Finally, we also checked proliferation at 28 dpa, when the redifferentiation phase is completed, and found very few positive cells both in WT and mutant digit tips (Fig. 5H-I’) ruling out the possibility of a delayed regeneration response in the mutant.

The above results identified several typical cellular and molecular blastemal features that in combination support the initiation of a regenerative response with the formation of a blastema in the amputated stump of *ΔLARM1/2* digits, though it failed to progress to regenerate the missing part.

### Comparison of the transcriptomic signatures of WT and mutant blastemas

Next, to uncover the factors responsible for the failed regeneration, we performed RNA-seq to compare the transcriptional signature of the regenerative (WT) versus the non-regenerative (*ΔLARM1/2* mutant) blastema. To this end, we collected WT and mutant digit tips including the blastema and overlying epidermis at 12 dpa and 14 dpa (Fig. 6A). Two WT and three mutant biological replicates were collected per stage, each one pooling together 6 digit tips. These two stages were selected based on our time-course experiment evaluating blastemal progression (Supplemental Fig. 2), to capture its maximum development (12 dpa), and the starting of blastemal differentiation (14 dpa).

**Figure 6.**
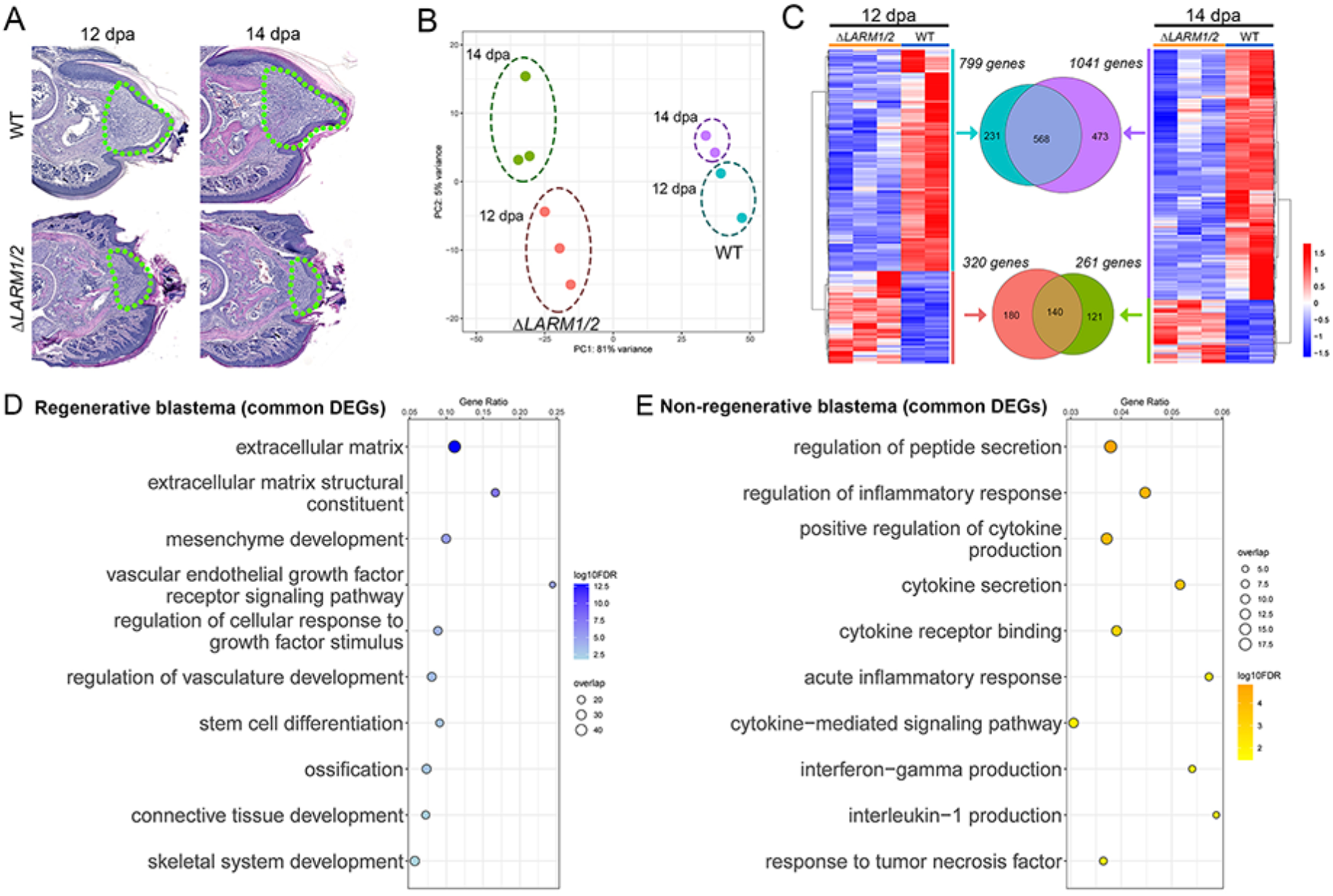
Transcriptomic comparison between WT regenerative and *ΔLARM1/2* non-regenerative blastemas. (A) Hematoxylin & eosin-stained longitudinal sections of amputated WT and *ΔLARM1/2* digit tips at the stages indicated on top. Green dotted lines outline the region dissected for the RNA-seq. (B) Principal component analysis (PCA) plot of the top 500 most variable genes, using normalized counts. Dashed circles depict same group conditions as indicated. (C) Sides:row-clustered heatmaps of row-scaled log2(1 + TPM) values of the DEGs obtained by Wald test of WT vs Δ*LARM1/2* at 12dpa (left) and 14dpa (right). Center: Venn diagrams showing common DEGs between both stages. (D-E) Dot plot of selected representative GO:BPs terms extracted from the functional annotation of the set of common DEGs upregulated in WT (D) and *ΔLARM1/2* (E) blastemas. X axis represents the Gene Ratio (proportion of DEGs in a given term). The size of the dots represents the number of DEGs in a given term and the color scale represents the negative value of Log10 of the adjusted p-value.

Libraries for RNA-seq were prepared from each of the 10 collected samples and sequenced with 50-bp paired-end read lengths resulting in an average of at least 80 million reads per sample from which nearly 86% of the reads mapped to unique loci. Only protein coding genes were used in the analysis.

When all samples were analyzed together, the principal component analysis (PCA) readily separated them by genotype (PC1 81% of the variance) (Fig. 6B). The second component, explained by 5% of the variance, segregated the samples by stage and revealed the proximity between the 12dpa and 14dpa blastema stages exposing the gradual progression through stages in both (WT and mutant) conditions. The control replicates were closer together than the mutant replicates perhaps reflecting a more canalized temporal course.

To gain insight into the gene regulatory networks involved in regeneration, we performed a differential expression analysis at each stage between WT and mutant samples using DESeq2 separately at each stage analyzed (Fig. 6C). At 12 dpa, we obtained a total of 1019 differentially expressed genes (DEGs) (log2FoldChange (L2FC) > 1.5 and p-adjusted value < 0.05), of which 799 were upregulated in the WT and 320 in the mutant (Table S1). Similarly, at 14 dpa, 1302 genes were differentially expressed of which 1041 were upregulated in the WT and 261 in the mutant (Table S2). Given that 74% of the differentially expressed transcripts were upregulated in the WT blastema that shows natural regenerative potential, it is possible that the main reason of the regeneration failure in the mutant is an inability to activate the appropriate transcription. We also reasoned that the transcripts upregulated in the mutant non-regenerative blastema, although much less in number, may provide valuable insights regarding the mechanisms that prevent or disrupt digit tip regeneration.

Having the list of DEG, we first looked at the expression of *Lmx1b* that, as expected, was elevated in WT (L2FC = 4.1, p-adj. value = 6.13e-16), by inspecting the coverage from RNA-seq across the *Lmx1b* locus (Supplemental Fig. 3A). *Lmx1b* expression, at very low level, was only detectable in the WT blastema, both at 12 and 14 dpa, but not in the mutant blastema. Because *Lmx1b* expression has not been previously assessed during digit tip regeneration, we used HCR RNA-FISH to validate these results and to determine the spatial distribution of *Lmx1b* transcripts (Supplemental Fig. 3B-E). During amphibian limb regeneration, *Lmx1b* is reactivated in the dorsal aspect of the blastema recapitulating the normal limb development pattern (Matsuda et al. 2001; Shimokawa et al. 2013; Satoh and Makanae 2014; Yamamoto et al. 2022). During digit tip regeneration, HCR detected a low level of *Lmx1b* expression in the WT regenerating blastema but, remarkably, with no DV bias (12 dpa; Supplementary Fig. 3B). Quantification of the signal by counting HCR RNA-FISH puncta in a standardized square area of the dorsal, ventral, proximal and distal blastema failed to detect expression differences. In agreement with the RNA-seq tracks, no *Lmx1b* transcripts were detected in the mutant blastema (12 dpa; Supplementary Fig. 3C-E). The uniform expression of *Lmx1b* throughout the WT blastema suggests that, if it plays a role during digit regeneration, it does not rely on asymmetric DV expression.

Next, we found that genes with reported expression in blastema and in digit tip regeneration were also expressed in the mutant blastema, although there was a general upregulation in the WT (Supplementary Fig. 4A). Examples of these genes include *Ltbp2, Arsi, Mest* (Storer et al. 2020; Johnson, Masias, and Lehoczky 2020), which were significantly upregulated in the WT, and *Cxcr4, Cxcl12* or *Dlx5* (Lee et al. 2013; Storer et al. 2020) which showed no significant differences. This observation furthers supports the presence of a blastema in the *ΔLARM1/2* stump.

To help understanding the mechanisms behind the different regenerative potential between WT and mutant blastemas, we functionally annotated the list of genes that were found upregulated in the WT blastema both at 12 dpa and 14 dpa (568 common DEGs, Fig. 6C and Table S3) and found significant enrichment (p-adjusted value < 0.05) for biological process (GO:BP) terms such as “ossification”, “collagen synthesis”, “mesenchymal development”, “ECM organization”, and other developmental processes (Fig. 6D and Table S4) all of them naturally occurring during regeneration. Also, there was a great enrichment of terms associated with the development of the vasculature and “vascular endothelial growth factor signaling pathway” (VEGF signaling pathway; Fig. 6D and Table S4) We also functionally annotated the list of genes upregulated in the non-regenerative mutant blastema both at 12 dpa and 14 dpa (140 common DEGs). With this set, the most enriched GO:BP terms were associated with inflammation and immune response including “regulation of inflammatory response” and “cytokines and inflammatory response” (Fig. 6E and Table S5). Given the importance of vascularization, ECM remodeling and inflammation during regeneration, we sought for further validation and investigation.

### Poor vascularization and aberrant ECM remodeling in the mutant blastema

Genes associated with vascularization and the VEGF signaling pathway are highly enriched in the WT blastema (Fig. 6D, Supplementary Fig 4B and Table S4) and may indicate that past an initial hypoxic phase a second phase of appropriate vascularization is also crucial for regeneration. It is possible that vascularization provides humoral trophic factors that foster regeneration. To explore the vascularization state of the mutant blastema compared to that of the WT control we analyzed the expression of *Cdh5*, a marker of endothelial cells, by ISH in tissue sections. *Cdh5* marked the presence of blood vessels that were conspicuous both in WT and mutant blastemas, although *Cdh5* transcription by RNA-seq was significantly upregulated in WT mice (Fig. 7A-A’ and Supplemental Fig. 4B). Transcripts of *Vegfa*, a growth factor essential for the proliferation and migration of vascular endothelial cells during angiogenesis and with other reported functions such as M2 macrophage polarization (Linde et al. 2012), were abundant in the WT blastema but very scarce in the mutant (Fig. 7B-B’).

**Figure 7.**
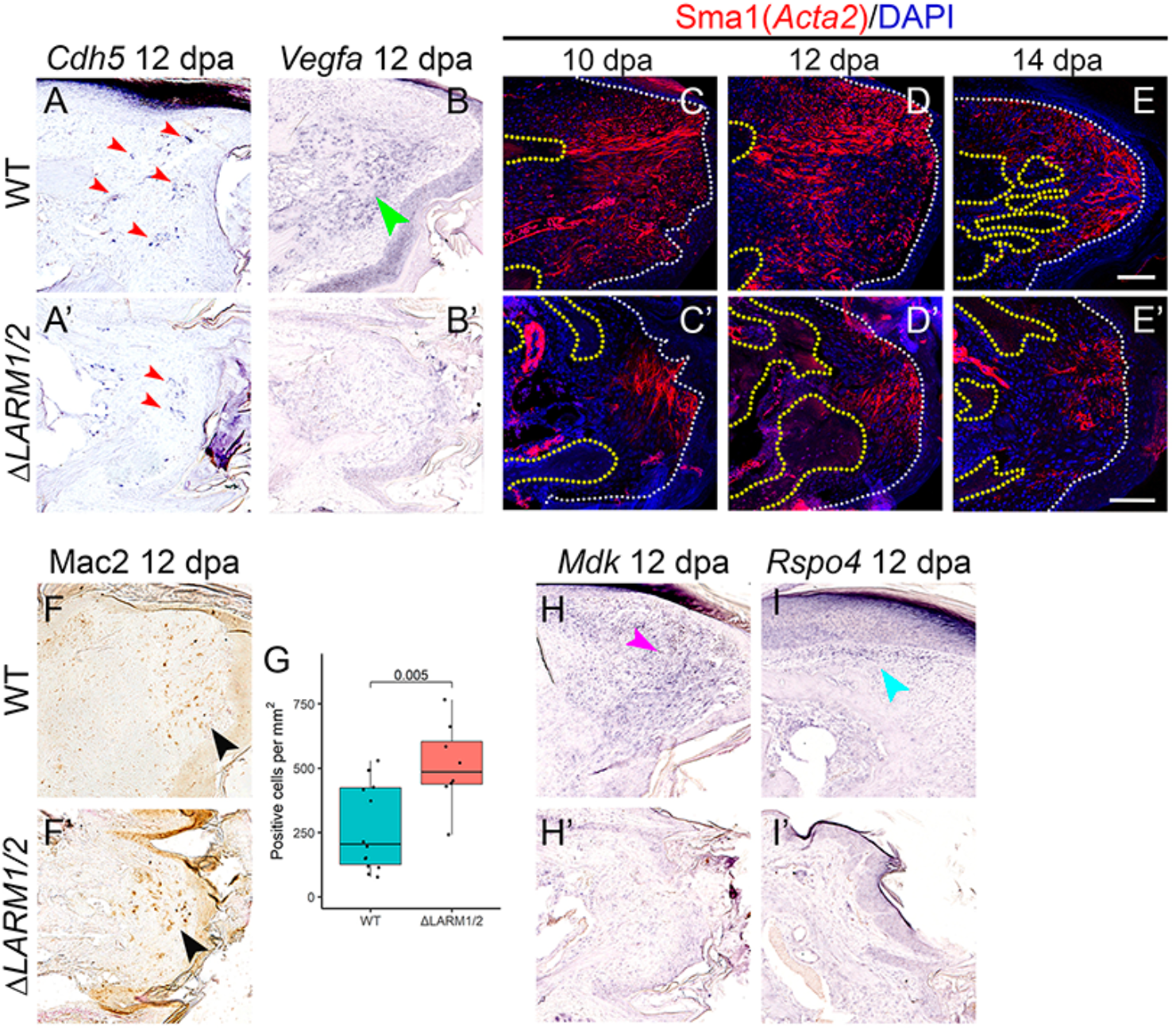
Histological validation of candidates extracted from RNAseq analyses reveals novel features in digit tip regeneration. (A,B’) ISH targeting *Cdh5* (A, A’) and *Vegfa* (B,B’, n=3) mRNA in 12 dpa (P33) longitudinal sections of WT (A,B) and *ΔLARM1/2* (A’,B’) digit tips. Red arrowheads depict regions of *Cdh5* expression corresponding to blood vessels. Green arrowhead depicts detected expression of *Vegfa*. (C-E’) IF for Sma antigen (red; encoded by *Acta2* gene, n=2) at the indicated stages in WT (C,D,E) and *ΔLARM1/2* (C’,D’,E’) blastemas. Blue corresponds to DAPI staining. White and yellow pointed lines outline epidermis and bone, respectively. (F,F’) IHC targeting Mac2 antigen (encoded by *Lgals3* gene, n=4) in 12 dpa (P33) longitudinal sections of WT (F) and *ΔLARM1/2* (F’) blastemas. (G) Boxplots representing the quantification of the number of Mac2 positive cells per mm^2^ in WT vs *ΔLARM1/2* blastema sections, with p-value calculated by Student’s t.test represented at the top. Each dot is a section quantified. n=4 biological replicates. (H,I’) ISH targeting *Mdk* (n=3) and *Rspo4* (n=3) mRNA in 12 dpa (P33) longitudinal sections of WT (H, I) and *ΔLARM1/2* (H’, I’) blastemas. Purple and teal arrowheads depict region of expression.

We also visualized the expression of the vascular smooth muscle marker Acta2 by IF using the anti-alpha smooth muscle actin antibody (Sma). Acta2/Sma was increased in the WT blastema when compared to the mutant blastema (Fig. 7C-E and Fig. 7C’-E’ and Supplemental Fig. 4B). Besides vascular pericytes and vascular smooth muscle cells, *Acta2* is also expressed in myofibroblasts, important players in wound healing that generate the needed contractile forces. Sma positivity revealed a strikingly different organizational pattern in mutant and WT blastemas. While in the WT, Sma positive fibrils were found all over the blastema with a predominant proximo-distal orientation, in *ΔLARM1/2* mutants the Sma positive fibrils concentrated under the wound epidermis mainly in a parallel orientation, suggesting the initiation of a fibrotic reaction. Collectively, the data show that the process of angiogenesis and ECM organization is altered in the mutant supporting the notion that fine modulation of revascularization at later stages is important for successful regeneration.

### Persistent inflammation in the mutant blastema

The amputation of the digit tip is followed by an early phase of inflammation and tissue histolysis with ECM remodeling that is essential for tissue healing and blastema formation. Then, the inflammation regresses and regeneration proceeds, as sustained inflammation may lead to regeneration failure (Gawriluk et al. 2020; Xu et al. 2019). This may be the case in the mutant blastema as we identified a number of genes related with inflammatory processes upregulated in the mutant blastema (Supplemental Fig 4C).

Because macrophages have been shown to be a critical cell type in the process, we investigated their presence by IHC using the anti Mac-2 (also known as Galectin3) antibody. Galectin3, encoded by the gene *Lgals3*, is a general marker of macrophages that is significantly upregulated in the mutant blastema (Supplemental Fig. 4C). The anti Mac-2 antibody detected a slight, but significant increase in the macrophage population in the mutant blastema compared to control (at 12 dpa, Fig. 7F-F’, G). Subpopulations of M1 and M2 macrophages are the main sources of pro-inflammatory and anti-inflammatory signals, respectively, and activated M2 macrophage phenotypes have been suggested to be involved in successful regenerative responses (Aztekin et al. 2020; Bijarchian et al. 2021; reviewed in Aztekin and Storer 2022). Remarkably, classical markers of pro-inflammatory macrophages (M1) including *TNFalpha, Il1b, Cxcl2* and *Csf2* (De Oliveira, Rosowski, and Huttenlocher 2016) were upregulated in the mutant blastema, while markers of M2 macrophages, such as Cd206 (*Mrc1*) and *Cx3cr1* (Burgess et al. 2019) were upregulated in the WT blastema (Supplemental Fig. 4C and D). These data support the presence of M2 macrophages in regenerative WT blastema and enrichment of pro-inflammatory M1 macrophages in the mutant blastema. The study of macrophage polarization/heterogeneity and the impact in digit regeneration deserves further investigation.

Along this line, Midkine (Mdk), a secreted growth factor strongly induced during inflammation, was recently shown to play a role both in the healing of wound epidermis and in the resolution of the initial inflammation in axolotl limb regeneration (Tsai, Baselga-Garriga, and Melton 2020). Interestingly, *Mdk* is upregulated in the WT regenerative blastema (L2FC = 1.73, p-adj value = 6,56e-8) and, accordingly, *Mdk* transcripts in the WT were abundant at the distal periphery of the blastema but undetectable in the mutant blastema (Fig. 7H-H’) indicating that it may play similar functions during digit regeneration. Other genes with reported anti-inflammatory functions such as *Lgals1* and *Siglec1* (Fernandez-Perez et al. 2021; Zheng et al. 2015) were also upregulated in the WT blastema (Supplemental Fig. 4E). Together, our data suggest that an abnormal inflammatory state in the mutant blastema may preclude the transition to a regenerative state.

Conjointly, the data also indicate that in the absence of *Lmx1b*, digit tip regeneration is halted beyond the formation of a small blastema in which inflammatory gene expression persists, re-vascularization is reduced, and ECM aberrantly remodeled or showing a fibrotic response.

#### Lmx1b may have a direct effect driving regeneration

The abnormal transcriptional signature of the *ΔLARM1/2* blastema may reflect an altered environment probably because of the lack of appropriate signaling from the nail-like DDP. Also, it is important to note that the position of the DDP is not completely equivalent to that of the nail in control digit tips (see Fig. 1E-E’). Thus, while the normal nail in its dorsiflexed position overlies the entire distal phalanx, the DDP overlies the proximal, but not the distal aspect of the straight double ventral distal phalanx.

However, the fact that *Msx1* expression is specifically lost in its dorsal domain, coincident with the region of *Lmx1b* expression, but not in the lateral domain (Fig. 2 F-H and Supplementary Fig. 5), led to the hypothesis that *Lmx1b* may be more directly involved in regenerative capacity through the establishment of a specific transcriptional state of the dorsal dermis that promotes blastemal progression. Correspondingly, during development this central dorsal *Msx1* expression, the part of the domain that overlaps with *Lmx1b* expression, is lost in *ΔLARM1/2* mutants suggesting direct Lmx1b regulation of *Msx1* (Supplementary Fig. 5). Interestingly, recent scRNA-seq analyses have identified a distinct transcriptional signature specific of this dermal population, different from that of the rest of mesenchyme (Storer et al. 2020; Kim et al. 2021). To get some insights into the transcriptional state of the mutant dorsal dermis, we explored in our DEGs the expression of characteristic genes such as *Rspo4, Sfrp2*, and *Crabp1* and all of them were found downregulated in the mutant blastema (Supplemental Fig. 4F; Han et al. 2008; Storer et al. 2020; Kim et al. 2021). Using ISH we validated *Rspo4*, an activator of the canonical Wnt signaling pathway, normally expressed along the nail dermis in the WT but that was undetectable in the mutant digit (Fig. 7I-I’). These results indicate that *ΔLARM1/2* digits lack the distinct dorsal dermis transcriptional signature that, together with the lack of *Lmx1b* expression in the blastema, points to a possible direct requirement of *Lmx1b* in regeneration, a hypothesis that requires further investigation.

## DISCUSSION

### Digits with no *Lmx1b* expression fail to regenerate

Digit tip regeneration offers a unique opportunity to study multi-tissue regeneration in mammals. Even considering that only a small proportion of the limb regenerates, the tip of the distal phalanx, it is worth studying because of the broad therapeutic potential for regenerative wound healing. The initial reports on digit tip regeneration both in humans and mice already identified a correlation between the nail and regeneration, as only amputations that preserved the nail matrix regenerated (Zhao and Neufeld 1995; Douglas, 1972). Since then, several studies have tried to understand this correlation although a direct test of the nail requirement for digit tip regeneration has not been possible because of the lack of appropriate models (Takeo et al. 2013; Lehoczky and Tabin 2015).

The nail is a dorsal appendage, the formation of which depends on correct dorsal-ventral patterning directed by Lmx1b, the dorsal limb determinant. *Lmx1b*-null mutants display double ventral digits with no nail but die perinatally because of other systemic Lmx1b-associated defects (Chen et al. 1998). However, in the absence of two limb-specific *Lmx1b* enhancers (*ΔLARM1/2* mutants), *Lmx1b-null* phenotype is recapitulated only in the limbs of mutants that are otherwise viable and similar to wild-type (Haro et al. 2021). The double-ventral digits of *ΔLARM1/2* mutants offer an extraordinary opportunity to test of the nail involvement in digit tip regeneration.

Interestingly, a detailed analysis of the digit tip of *ΔLARM1/2* mutants revealed that the dorsal digital pad (DDP) that formed, in lieu of the nail, progressively developed some nail-specific epidermal characteristics including expression of *Lgr6* and *Hoxc13* considered markers of nail stem cells (Lehoczky and Tabin 2015; Fernandez-Guerrero et al. 2020). Concomitantly, this dorsal pad also contained eccrine glands normally only associated with volar pads. This reflects the hybrid nail-pad nature of the DDP and reveals that some dorsal characteristics of the epidermis can occur independently of the normal *Lmx1b*-dependent mesenchymal interaction. It should be stressed that the *ΔLARM1/2* mutant digits showed no dorsal musculoskeletal features, i.e., the bones, tendons and other elements adopted a double ventral organization. Equally, the *Msx1* expression characteristic of the nail associated dermis was not detected.

Despite the presence of this hybrid nail-pad structure, we demonstrate in this report that *ΔLARM1/2* mutants lack digit tip regeneration supporting the concept that a well-developed nail organ is a prerequisite for regeneration.

### Digit tip regeneration is possible in the absence of DV polarity

In salamander, the absence of anterior-posterior polarity cues abrogates limb regeneration, but when positional disparities are restored regeneration occurs (Tanaka 2016). Although conflicting reports on the ability of double-dorsal or double-ventral limbs to regenerate have been reported (Bryant and Gardiner 2016; Burton, Holder, and Jesani 1986), we considered the alternative possibility that DV polarity was required for digit tip regeneration. To test this, we used the recently described *Del(27)* mutant generated from the deletion of *Maenli* lncRNA that regulates the limb-specific expression of *En1* (Allou et al. 2021). In the absence of limb-specific *En1*, ectodermal *Wnt7a* and mesodermal *Lmx1b* extend to the ventral limb and form double dorsal digits bearing circumferential or conical nails with no DV polarity. Here we show that *Del(27)* mutants regenerate normally demonstrating that disparities in DV positional information are dispensable for digit tip regeneration.

### In the absence of *Lmx1b*, a defective blastema forms after digit tip amputation

The amputation of the digit tip initiates a complex series of events including inflammation, tissue histolysis, and ECM remodeling that leads to delayed wound closure (compared to normal, scar-related wound healing), and the subsequent formation of the blastema. In the absence of *Lmx1b* expression, the regeneration process is initiated normally, with histolysis and wound closure occurring similar to the WT, but the process stalls after the formation of a blastema. This is reminiscent of the situation in *Xenopus* froglets (Suzuki et al. 2006), in the accessory limb model (Endo, Bryant, and Gardiner 2004) and in proximal non-regenerative amputations of the digit tip (Dawson et al. 2020), situations in which failed regeneration still associates with the formation of a blastema-like structure.

The presence of the blastema in the non-regenerative *ΔLARM1/2* digits is confirmed by several characteristic cellular and molecular features including cell-cycle reactivation and nerve colonization. However, markers of redifferentiation such as the osteogenic markers were not detected in the mutant blastema indicating an absence of progression factors or an inability to respond to them and differentiate.

To identify the molecules and pathways that render the mutant blastema regeneration incompetent, we compared its transcriptional signature with that of the control blastema, at two different stages and found the expression of genes associated with vascularization and ECM organization to be enriched in the control, while the expression of genes associated with inflammation were enriched in the mutant.

It is known that the digit tip blastema is avascular in the initial stages up to stages close to starting redifferentiation. The hypoxia of the initial blastema is critical for a successful regenerative response and when angiogenesis is precociously introduced, regeneration is inhibited (Yu et al. 2014; Sammarco et al. 2015). Vascularization of the blastema occurs in the redifferentiation phase when bone formation is initiated (Fernando et al. 2011). We show that the deficient expression of angiogenic genes in the mutant blastema correlated with a poor invasion of blood vessels suggesting that angiogenesis needs to be tightly modulated throughout the regeneration process for a successful regeneration outcome, possibly through the supply of humoral factors. Prevention of angiogenesis in early blastema stages depends on the upregulation of the anti-angiogenic factor *SerpinF1/Pedf* (Becerra and Notario 2013; Yu et al. 2014). Curiously, *Pedf* is downregulated in the mutant blastema suggesting that a persistent block in vascularization is not the cause of the poor vascularization of the mutant blastema.

The ECM also plays a key role in regulating the injury response by recruiting and organizing progenitor cells (Marrero et al. 2017). We show that in the WT blastema Sma positive cells arrange longitudinally along the proximo-distal axis, from the bone stump to the wound epidermis, similar to the pattern of ECM fibers produced by fibroblast reticular cells (Marrero et al., 2017). This ECM organization in the regenerating blastema is in stark contrast with the ECM arrangement of the mutant blastema that has Sma positivity rather parallel to the overlying wound epithelium, consistent with scar formation. The myofibril connection between the tip of the stump and the wound epidermis in regenerating blastemas suggests a previously unnoticed mechanical interaction between these two structures.

In addition to these differences, our analysis also showed that the expression of inflammation-associated genes was enriched in the mutant blastema. During the early stages after amputation, macrophages play a critical role in limb (Godwin, Pinto, and Rosenthal 2013; Jennifer Simkin, Sammarco, et al. 2017) and other regenerative scenarios such as the zebrafish fish fin (Petrie et al. 2014) and the ear hole punches in Spiny mice (Jennifer Simkin, Gawriluk, et al. 2017). At the stages analyzed in our study, we detected a significant increase in macrophage numbers in the mutant blastema while markers of M2 polarization and genes with reported anti-inflammatory functions are found upregulated in the WT blastema. Together, our results revealed that while inflammation is resolved with the progression to redifferentiation in the WT blastema, in the *Lmx1b*-deficient blastema inflammation persists and regeneration stalls.

### Is *Lmx1b* necessary for digit tip regeneration?

The failure of *ΔLARM1/2* mutants to regenerate may be attributed, at least in part, to an insufficient signaling from the DDP nail-like epithelium. The presence of nail stem cells and Wnt signaling in the DDP epidermis may not be sufficient and/or may be positioned too proximal for adequate signaling. However, the absence of a molecularly competent dorsal dermis in the mutant could potentially support the alternative scenario of an onychodermis unable to convert the epidermal signals into a regenerative response.

The dorsal dermis is characterized by the expression of the transcription factor Msx1 and mice null for *Msx1* are unable to regenerate (Allan et al., 2006; Han et al., 2003; Han et al., 2008). *Msx1* is not detected in the blastema suggesting that it functions in the dorsal dermis (Han et al., 2008; Lechoczky et al., 2011). Here, we show that in *ΔLARM1/2* mutants the expression of *Msx1* is specifically lost in a dorsal domain coincident with the area in which *Lmx1b* is expressed in WT digit tips. Therefore, it seems reasonable to speculate that it is the absence of *Lmx1b* that leads to absence of *Msx1* and therefore to an incompetent dorsal dermis. This, together with the low and uniform *Lmx1b* expression throughout the WT regenerative blastema but no in the mutant blastema, as examined by HCR RNA-FISH and bulk mRNA-seq data, suggest that *Lmx1b* may be an important key factor driving regeneration. It is important to note here that the lack of *Lmx1b* expression in the mutant blastema rules out the possibility of *Lmx1b* reactivation through the use of regeneration-specific enhancers.

The origin of the blastema cells has not been completely determined but no change in location of grafted cells has been reported suggesting no considerable/major cell rearrangements (Wu et al. 2013). Thus, the uniform *Lmx1b* expression in the blastema suggests *de novo* activation at least in those blastema cells that do not derive from *Lmx1b* expressing cells in the dorsal dermis. *Lmx1b* responds to Wnt signaling and could be activated by the nail dependent Wnt environment.

In summary, we show that double ventral digits with no *Lmx1b* expression do not elicit a regenerative response, but that very likely this is not due to the lack of DV polarity as double-dorsal digits do regenerate. Similar findings have been recently reported using conditional knockouts of *Lmx1b* and *En1* (Johnson et al. 2022). Even considering that the lack of a fully developed nail organ might impair the regenerative outcome, our results suggest that the expression of *Lmx1b*, which normally persists in the connective tissue underlying the nail through adulthood, may be required in the onychodermis to promote blastemal progression to regenerate the digit tip. The functional implication of *Lmx1b* expression in the blastema merits further investigation.

## Acknowledgment

This study was supported by funds from the Pathology Research endowment and the Crook’s Chair to Kerby C. Oberg and the Spanish Ministry of Science and Innovation Grant PID2020-114525GB-I00 to Marian Ros. Alejandro Castilla-Ibeas is supported by Spanish Ministry of Science and Innovation PhD fellowship FSE/AEI/PRE2018-083421. Sofía Zdral was supported by a PhD fellowship from the University of Cantabria. We thank Can Aztekin for critical reading and the Animal Facility of the University of Cantabria for outstanding animal husbandry.

---

**Figure S1, related to Figure 4**

**P3 *ΔLARM1/2* mutants do not regenerate amputated digit tips.**

(A-A’) Masson’s trichrome staining of longitudinal sections of WT (A) and *ΔLARM1/2* mutant (A’) P3 digit tips. The plane of amputation is depicted by a dashed line (n=4).

(B-C’) μCT renderings of WT (B-C) and *ΔLARM1/2* mutant (B’-C’) unamputated contralateral digit tips (P24, B-B’) used as controls of P3 amputated digits at 21 dpa (C-C’) (n=4).

(D-D’) Masson’s trichrome staining of longitudinal sections of WT (D) and *ΔLARM1/2* mutant (D’) P3 amputated digits at 21 dpa (P24) (n=4).

Scale bars: 500 μm.

**Figure S2, related to Figure 5**

**Histologic time course of digit tip regeneration**

(A-E’) Masson’s trichrome staining of WT (A-E) and *ΔLARM1/2* mutant (A’-E’) digits amputated at P21 and analyzed at the stages indicated on top.

Scale bars: 500 μm

**Figure S3, related to Figure 6**

**Low levels of unpolarized *Lmx1b* expression in WT, but not *ΔLARM1/2* blastemas.**

(A) *Lmx1b* coverage plot from the blastema RNA-seq experiments, RNA-seq sample is indicated at the right.

(B) Left, schematic of the 12 dpa blastema digit tip depicting *Lmx1b* detected expression. Dashed boxes indicate the regions quantified in using HCR-FISH. Right, boxplot representing number of puncta in each region indicated, analyzed by HCR-FISH. Statistical significance was assessed by ANOVA (n=3).

(C, D) HCR-FISH representative images of the quantification performed in 12 dpa (P33) blastema of WT (D) and *ΔLARM1/2* (E) digit tips. Scale bar: 20 μm

(E) Boxplot representing number of puncta detected in the 12 dpa WT and Δ*LARM1/2* blastemas. Student’s T-test was used to assess the statistical significance (n=3).

**Figure S4, related to Figure 6**

**Analysis of DEGs grouped in digit tip regeneration-related processes.**

(A-F) Scaled heatmaps of Log2(1+TPM) for the indicated genes, grouped by processes as indicated at the top of each one. Asterisks denote genes that were validated. In A, nonsignificant (ns) genes are indicated on the left.

**Figure S5, related to Figure 6.**

A distinct dorsal *Msx1* expression domain is specifically lost in the absence of *Lmx1b*

(A-A’) mRNA ISH for *Msx1* and (B-B’) HCR RNA-FISH for *Lmx1b* in transverse sections E15.5 digit tips of WT (A, B) and *ΔLARM1/2* mutants (A’, B’). In the WT, *Msx1* transcripts are found in the dorsal and lateral mesoderm while the dorsal domain, coincident with the expression of *Lmx1b* is lost in the mutant.

(C-C’) Corresponding schematics representing the expression domains of *Msx1* and *Lmx1b*

## Material and Methods

### Mouse strains and animal ethics

All animal procedures were designed and performed according to the European Union regulations and 3R principles. These were reviewed and approved by the Bioethics Committee of the University of Cantabria. We have used the previously published Δ*LARM1/2* and the *Del(27)* mutant lines generated by CRISPR/Cas9 (Haro et al. 2021; Allou et al. 2021). A minimum of 2 independent specimens were analyzed for each experiment and time point analyzed.

### Surgical procedures

Digit amputations were carried out on postnatal day 3 (P3) and on P21 mice as previously described (J. Simkin et al. 2013). Briefly, the distal third of the distal phalanx of the three central hindlimb digits was surgically removed using a #10 disposable scalpel (P21 WT) or a microsurgery scissor (P3 and P21 *ΔLARM1/2* mutants) (Ref: 14060-10, F.S.T. tools). Three central contralateral digits remained untouched as controls, except from mice utilized for RNA-seq experiments, whose contralateral central digits were also amputated and harvested at the indicated timepoints to ensure viability of the experiment.

### Histological Analysis

Dissected tissues were fixed in 4% paraformaldehyde for 48 hours at 4°C, decalcified in EDTA 9% (pH=7.4) for a variable period of time according to age, paraffin embedded and sectioned (7 μm) in a Leica microtome. Hematoxylin–eosin staining and Masson’s trichrome, were performed following standard procedures.

### Immunohistochemistry (IHC) and immunofluorescence (IF)

IHC and IF were performed on 7 μm sections using standard procedures. Antigen retrieval, when required, was performed using the 2100 Antigen Retriever (www.antigen-retriever.com/). IHC signal was amplified by using biotin conjugated secondary antibodies combined with VECTASTAIN® Elite ABC-HRP Kit (Vector laboratories) and images acquired with a Zeiss Z1 scanner. For IF samples were incubated with Alexa488-conjugated anti-rabbit or anti-mouse secondary antibodies (1:500) and counterstained with DAPI (1 μg/μL) and images acquired either on a Zeiss Axio M1 epifluorescence microscope or on a Leica Sp5 confocal microscope at the green (Em: 470/40) and blue (Em: 445/50) channels respectively. Autofluorescence was removed by subtracting the signal from the red channel with the Fiji software. Primary antibodies used were: anti-Krt5 (Sigma, SAB4501651), anti-Krt10 (BioLegend, 905401), anti-Pan-Cytokeratin AE13 (Santa Cruz, sc-57012), anti-Krt8/18 (DSHB, AB_531826), anti-Mac2 (CEDARLANE, CL8942B), anti-Actin (SMA; Sigma, C6198), anti-Ki67 (Abcam, Ab1667), anti-phospho-histone H3 (Upstate, 06-570), and anti-TUBB3 (Tuj1; BioLegend, 811801). All antibodies were used at 1:1000 concentration.

### Micro computed tomography (μCT)

Digit tips were arranged with some adaptation to what is described in Hipsley et al. 2020 which allows scanning of hundreds of samples in the same container. Then the preparations were scanned with Skyscan1172 at 40 kV, 100 μA, and 27.03 μm pixel resolution. Subsequent reconstruction was performed using the NRecon reconstruction software (Bruker) and visualized and imaged with CTvox software (Bruker).

### In situ hybridization (ISH) and Hybridization Chain Reaction (HCR) - Fluorescence In Situ Hybridization (FISH)

ISH was performed with RNA digoxigenin-labeled probes following standard procedures. with WT and mutant samples processed in identical conditions. The primers used to amplify cDNA templates for riboprobe synthesis are included in key resources table. For HCR, FFPE tissue protocol qHCR v3.0 (Molecular Instruments) was followed, with adaptations. Namely, 7 μm sections were deparaffinized and subjected to 10 minutes proteinase K treatment at 37°C (10 μg/mL), hybridized overnight at 37°C with 40 probe pairs against *Lmx1b* coding sequence (NM_010725.3), amplified overnight with B3 amplifiers conjugated with Alexa-647 fluorophores, and counterstained with DAPI (1 μg/mL). Mosaic images of E11.5 hindlimbs and E15.5 and P3 digit tips were obtained using a 63X/1.4 objective and the confocal Leica SP5 microscope.

### HCR quantification

For *Lmx1b* quantification in P21 uninjured digit tips and 12 dpa blastemas, a minimum of 3 z-stacked images (12 μm depth, 2 μm step size) of each analyzed region from three biological replicates were obtained under a 100X objective with a Zeiss M1 epifluorescence microscope. Images were processed using Image J software. First, background noise was filtered out by a median filter and point contrast was enhanced using a Laplacian of Gaussian filter. Maximum projections of the stacked images were then obtained, and the number of points counted above the estimated threshold. Sections stained in parallel but without *Lmx1b* probes were used as negative controls to estimate point intensity threshold.

### Quantification of blastema area and number of Mac2 positive cells

Both were performed with QuPath software v0.3.0. Briefly, for blastema area quantification, 7 μm WT and *ΔLARM1/2* longitudinal sections of 12 dpa digit tips were analyzed by delineating the tissue distal to the bone stump that did not include the epidermis, and the area value exported for plotting and statistical analysis. For the Mac-2 cell quantification, positive cells were identified in longitudinal sections of 12 dpa WT and *ΔLARM1/2* digit tips by the automatic positive cell detection tool based on DAB intensity threshold, which was adjusted to reduce false positives. Positive cells per mm^2^ of blastema was calculated and exported for plotting and statistical analysis.

### RNA-seq

The three central digits of each hindlimb of four WT and six *ΔLARM1/2* mice were amputated as described previously. At 12 and 14 dpa equal number of mice were sacrificed and the 6 blastemas from each mouse were dissected from the area distal to the bone stump and pooled. RNA extraction was performed with QIAGEN RNA easy plus kit (QIAGEN) following manufacturer’s protocol. The quality and quantity of the RNA was assessed with a TapeStation Analyzer (Agilent). Illumina Stranded total RNA library preparation kit was used and the libraries sequenced with a NovaSeq6000 sequencer. Around 20 million 50-bp, paired-end reads per lane were obtained per sample and 4 lanes used for each sample.

### RNA-seq data processing

RNA-seq reads were mapped with STAR version 2.7.8a (Dobin et al. 2013) against the mouse reference genome (GRCm39) using the ENCODE parameters. Genes and isoforms were quantified with RSEM version 1.3.0 (Li and Dewey 2011) with default parameters using the gencode annotation version M27 with the primary assembly gtf (PRI). BigWig files were obtained from bam files and coverage plot was obtained using pyGenomeTracks package (Lopez-Delisle et al. 2021), with Lmx1b mm39 coordinates. Downstream analysis was performed with R (version 4.0.3; R core team, 2021).

### RNA-seq analysis

Counts and TPM were subset to protein coding genes and input in DESeq2 (Love, Huber, and Anders 2014, v1.30.1). Variance stabilizing transformation was used for PCA. DEGs were calculated by Wald test between 12 dpa and 14 dpa samples separately. The intersection of both DEGs list was obtained and functional annotation was performed with R package “WebGestaltR” (v0.4.4; Liao et al. 2019).

### Data availability

Raw data is deposited to GEO database under the accession number xxx (data under revision). All scripts used for this paper are available at Github repository https://github.com/AleCasIbe/scriptsforCastillaIbeasEtAl2022.

## Supplemental information

Document S1. Figures S1-S5

Table S1. List of differentially expressed genes in WT vs mutant samples at 12 dpa with statistical data and TPM for each sample, related to figure 6C.

Table S2. List of differentially expressed genes in WT vs mutant samples at 14 dpa with statistical data and TPM for each sample, related to figure 6C.

Table S3. Overlapping genes from the intersection of 12 dpa and 14 dpa lists of differentially expressed genes with statistical data and TPM for each sample, related to figure 6C.

Table S4. Functional annotation results from common DEGs at 12 and 14 dpa that were upregulated in the WT blastema, related to figure 6D.

Table S5. Functional annotation results from common DEGs at 12 and 14 dpa that were upregulated in the mutant blastema, related to figure 6E.

