## SUPPLEMENTAL TEXT AND FIGURES for "Failure of digit tip regeneration in the absence of *Lmx1b* suggests Lmx1b functions disparate from dorsoventral polarity"

Includes:

**Supplemental Figure 1**

**Supplemental Figure 2**

**Supplemental Figure 3**

**Supplemental Figure 4**

**Supplemental Figure 5**

**Supplemental Figure 1**

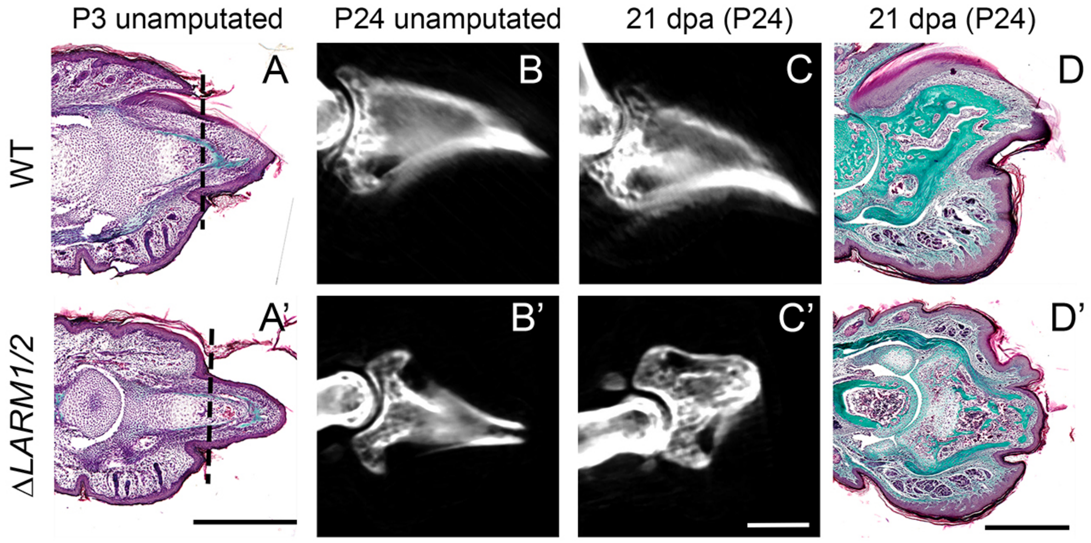


**P3 *ΔLARM1*/2 mutants do not regenerate amputated digit tips.**

(A-A') Masson's trichrome staining of longitudinal sections of WT and mutant P3 digit tips. The plane of amputation is depicted by a dashed line (n=2).
(B-C') µCT renderings of WT and mutant unamputated contralateral digit tips (P29) used as controls of P3 amputated digits at 21 dpa (C-C') (n=2).
(D-D') Masson's trichrome staining of longitudinal sections of WT and mutant P3 amputated digits at 21 dpa (P24) (n=2).
Scale bars: 500 µm.

**Supplemental Figure 2**

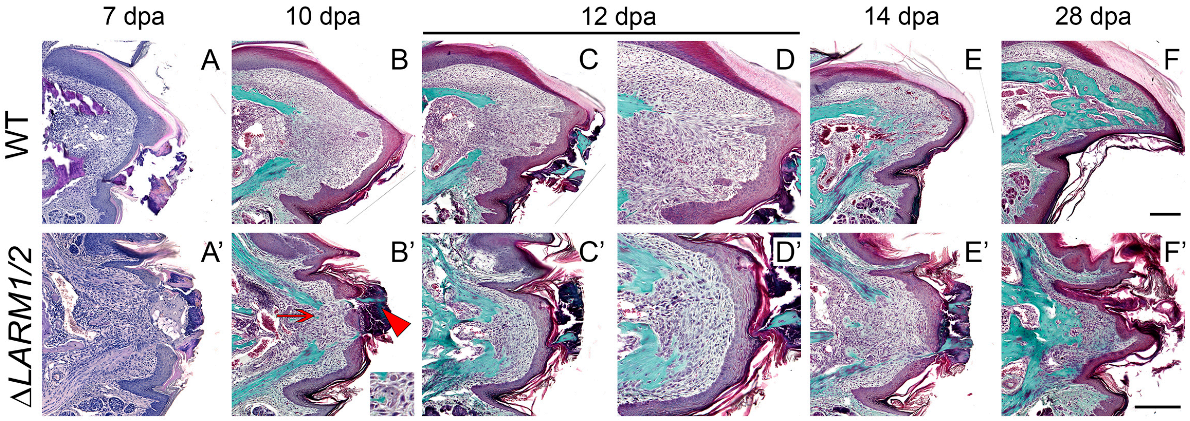


**Histologic time course of digit tip regeneration**

(A-E') Masson's trichrome staining of WT and Δ*LARM1/2* mutant digits amputated at P21 and analyzed at the stages indicated on top.

Scale bars: 500 µm

**Supplemental Figure 3**

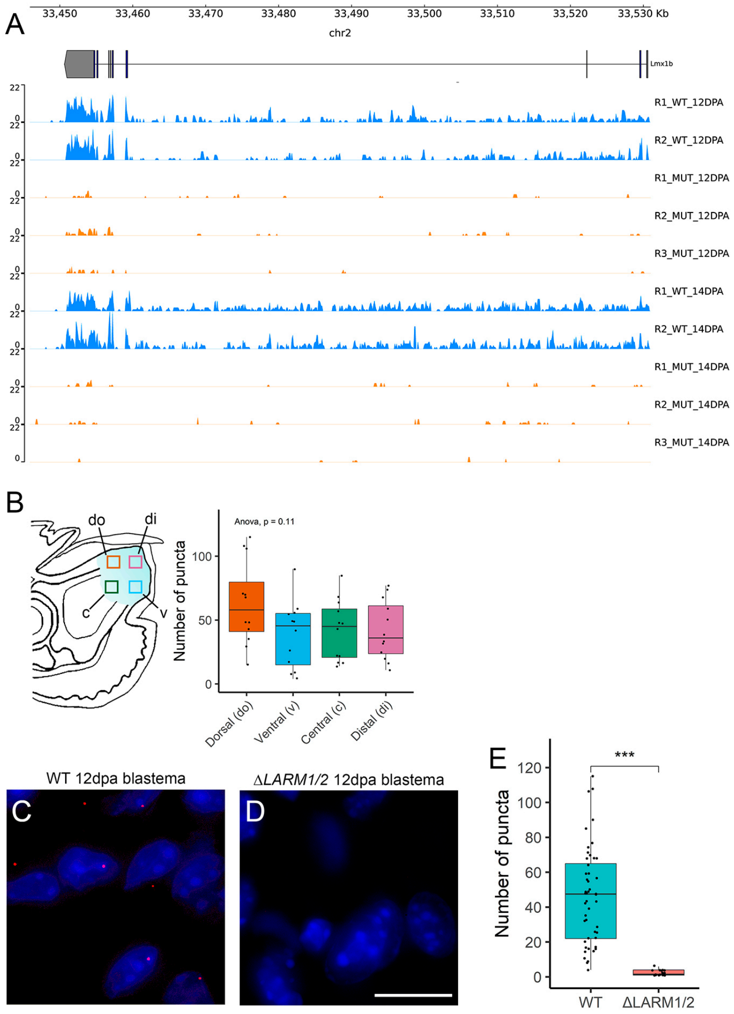


**Low levels of unpolarized *Lmx1b* expression in WT, but not Δ*LARM1/2* blastemas.**
(A) *Lmx1b* coverage plot from the RNAseq experiments, RNA-seq sample is indicated at the right.
(B) Left, schematic of the 12 dpa blastema digit tip depicting *Lmx1b* detected expression. Dashed boxes indicate the regions quantified in using HCR-FISH. Right, boxplot representing number of puncta in each region indicated, analyzed by HCR-FISH. Statistical significance was assessed by ANOVA (n=3).
(C, D) HCR-FISH representative images of the quantification performed in 12 dpa (P33) blastema of WT (D) and Δ*LARM1/2* (E) digit tips. Scale bar: 20 µm
(E) Boxplot representing number of puncta detected in the 12 dpa WT and Δ*LARM1/2* blastemas. Student's T-test was used to assess the statistical significance (n=3).

### Supplemental Figure 4

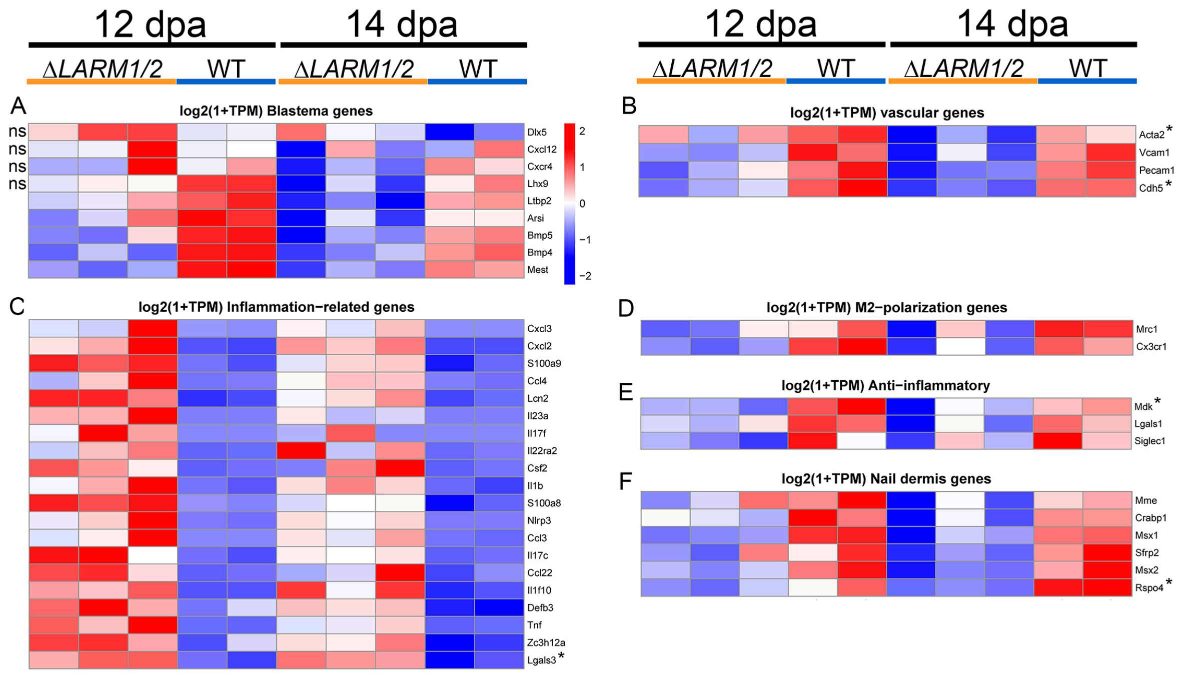


**Analysis of DEGs grouped in digit tip regeneration-related processes.**
(A-F) Scaled heatmaps of Log2(1+TPM) for the indicated genes, grouped by processes as indicated at the top of each one. Asterisks denote genes that were validated. In A, non-significant (ns) genes are indicated on the left.

**Supplemental Figure 5**

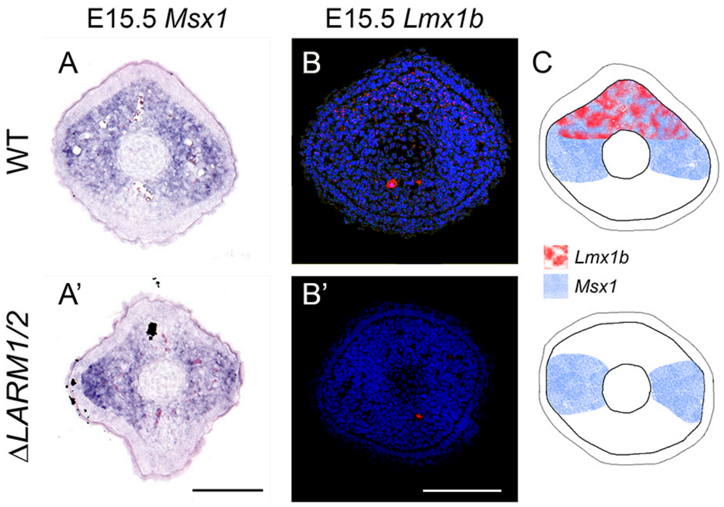


***Msx1* has a distinct dorsal expression domain that is specifically lost in the absence of *Lmx1b***

(A-A') mRNA ISH for *Msx1* and (B-B') HCR RNA-FISH for *Lmx1b* in transverse sections E15.5 digit tips of WT (A, B) and Δ*LARM1/2* mutants (A', B'). In the WT, *Msx1* transcripts are found in the dorsal and lateral mesoderm while the dorsal domain, coincident with the expression of *Lmx1b* is lost in the mutant.

(C-C') Corresponding schematics representing the expression domains of *Msx1* and *Lmx1b*
